# Genetic characteristics of *Apodemus speciosus* at Akiyoshidai Quasi-National Park in Yamaguchi Prefecture

**DOI:** 10.1101/2021.03.21.436340

**Authors:** Hiroyuki Imai, Hiroshi Tanaka, Taiki Matsuo, Miho Seto, Sumito Matsuya, Muneyoshi Hyoto, Kiyoshi Kano, Ken Takeshi Kusakabe

## Abstract

The large Japanese field mouse (*Apodemus speciosus*) is a small rodent endemic to Japan. The mice have a genetic characteristic in which the number of chromosomes differs between those from western Japan and those from eastern Japan. *A. spesiosus*, found throughout Japan, is used as a model animal for geogenetics and monitoring of radiation effects of wildlife. In this present study, to elucidate the genetic characteristics of the mice Akiyoshidai Quasi-National Park in Yamaguchi Prefecture, we investigated mitochondrial DNA and chromosome numbers. As a result, *A. speciosus* from Yamaguchi Prefecture were classified into the Honshu-Shikoku-Kyushu group and had a western Japan-type chromosome set of 2n=46; however, some Yamaguchi Prefecture mice formed a genetic cluster in Yamaguchi Prefecture, suggesting that continuous monitoring is needed to reveal the dynamics of genetic diversity.

## Introduction

The large Japanese field mouse (*Apodemus speciosus*) is a small rodent species endemic to Japan. *A. speciosus* inhabit the entire Japanese islands except for Okinawa and is frequently used as a model for studies of geographic isolation. The genetics of *A. speciosus* is characterized by different chromosome numbers in the east and west of Japan within a species. This characteristic karyotype is caused by a Robertsonian translocation (Shimba and Kobayashi 1969) and these translocated chromosomes were detected by FISH analysis (Yamagishi *et al.* 2012), which indicate that the mice are important species for genetic research.

Recently, *A. speciosus* was used as animals to monitor the effects of radiation around nuclear power plants. Especially, *A. speciosus* was used to monitor spermatogenesis and chromosomal abnormalities in Fukushima Prefecture (Okano *et al.* 2016; Takino *et al.* 2017; Ariyoshi *et al.* 2018; Fujishima *et al.* 2020). Decreases in the number of hematopoietic progenitor cells and chromosomal abnormalities were reported (Ariyoshi *et al.* 2020; Kawagoshi *et al.* 2017), indicating that *A. speciosus* was actually important in clarifying the effects of radiation on wildlife.

Although much genetic analysis has been performed, the sequence information is not sufficient to cover the whole of Japan. Many geogenetic studies were performed especially in the Japanese islands, but the information available on Honshu is not comprehensive. In the island genetics research, genetic diversity in the Seto inland sea region, Hokkaido and other remote islands were reported (Sato *et al.* 2017; Suzuki *et al.* 2015). The research leads to the elucidation of the genetic diversity of *A. speciosus* on the islands; however, *A. speciosus* needs more genetic consideration in each region of Honshu.

In this present study, we focused on mitochondrial DNA (mtDNA) sequences and chromosome numbers to clarify the genetic information of *A. speciosus* in Yamaguchi Prefecture, mainly in Akiyoshidai Quasi-National Park. Since Akiyoshidai is also designated as a special natural monument by the Japanese government, which usually restricts the collection of plants and animals, this present study could be an important record of natural history.

## Materials and Methods

### Study area and animals

Eight Japanese field mice were captured in Yamaguchi city and Akiyoshidai Quasi-National Park using Sherman trap (Fig. 1). The captures were performed with the permission of Yamaguchi Prefecture and Mine city. Captured mice were euthanized by cardiac blood sampling under isoflurane anesthesia. All the procedures using animals were approved by the Experimental Animal Care and Use Committee of Yamaguchi University (protocol number: 432).

**Fig. 1.**
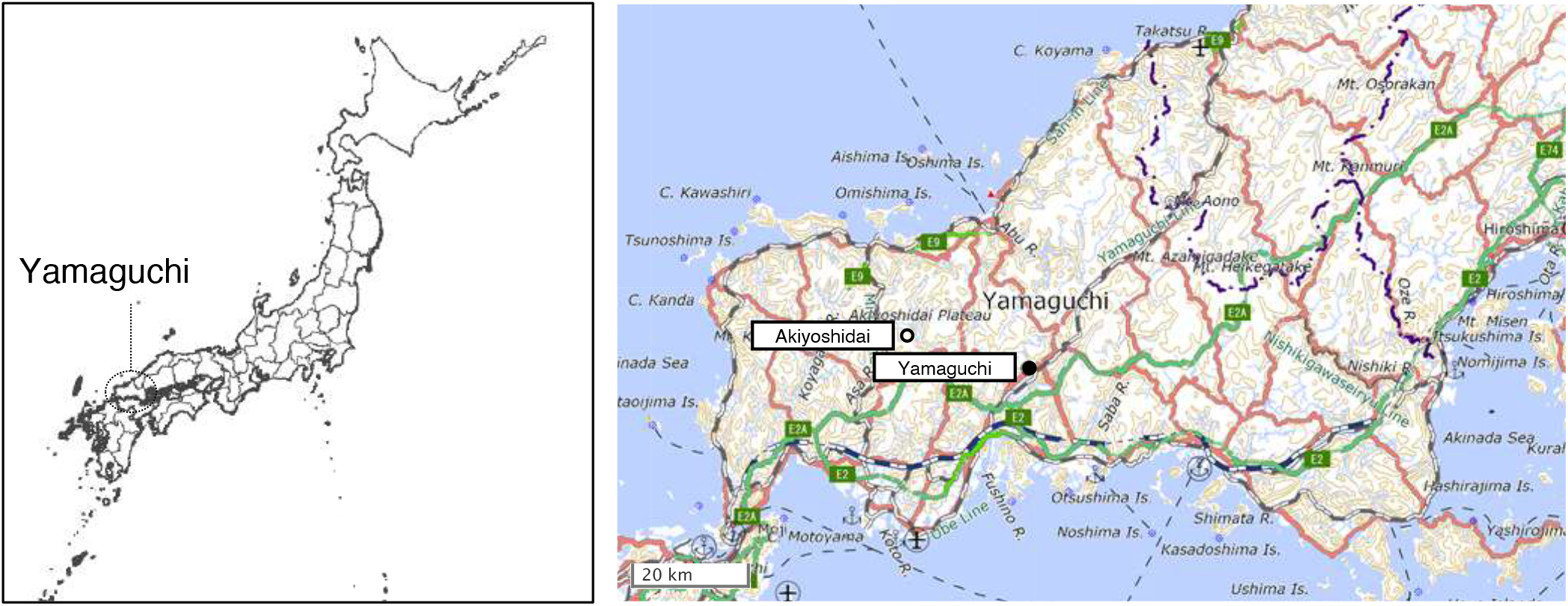
Sampling locations in this present study. The left map showed a general view of the Japanese archipelago and Yamaguchi Prefecture (dotted circle). The right map showed the locations of Yamaguchi city (filled circle) and Akiyoshidai (opened circle).

### DNA extraction and analysis

Genomic DNA was isolated from tail skin using NucleoSpin Tissue XS (Takara Bio, Shiga) according to manufacturer’s protocol. PCR was performed using PrimeSTAR HS (Takara Bio) and T100 Thermal Cycler (Bio-rad, CA). Primer sequences and their annealing temperatures were shown in Table S1. Fragments after electrophoresis were recovered using NucleoSpin Gel and PCR Clean-up (Takara Bio). The nucleotide sequences were determined by the Yamaguchi University Center for Gene Research. The obtained sequences were analyzed using ApE and MEGA X software (Tamura *et al.* 1993; Kumar *et al.* 2018; Stecher *et al.* 2020).

### Culture and chromosomal spread

The tail tips were placed on a 24-well plate and cultured in DMEM (Fujifilm-Wako, Tokyo) supplemented with fetal bovine serum (10%, Thermo Fisher Scientific Japan, Tokyo) and Penicillin-Streptomycin Solution (1x, Fujifilm-Wako). Passages and expansion cultures were performed using the media and Trypsin-EDTA solution (1 mmol/l EDTA-4Na, 0.25 w/v%, Fujifilm-Wako). The cells were arrested in metaphase by adding colchicine (Sigma-Aldrich Japan, Tokyo), suspended in hypotonic solution (1% sodium citrate), fixed in Carnoy’s fixation solution, and expanded onto glass slides. Chromosomes were stained with Giemsa Stain Solution (Fujifilm-Wako). A set of 30-50 chromosomes per individuals were observed.

## Result and Discussion

The Cytb and D-loop regions of mtDNA in large Japanese field mice (*Apodemus speciosus*) were analyzed using genomic PCR. The nucleotide sequences obtained by direct sequencing were deposited in the DDBJ (Table S2). Comparing the Cytb sequences of the captured mice in this present study with those in the database, the sequences of *A. speciosus* in Yamaguchi prefecture formed a cluster (Fig. 2, bold line). However, *A. speciosus* in Yamaguchi prefecture captured in this present study (Fig. 2 filled circles) and previous report (Fig. 2, opened circle; Suzuki *et al.* 2015) together were found to belong to the Honshu-Shikoku-Kyushu cluster. The distinct grouping of the mice in Hokkaido, Izu Islands, Sado Island, Nansei Islands and Tsushima Islands (Fig.2, fine, dotted, short-dashed, long-dashed and gray lines, respectively) might reflect previous report (Tsuchiya 1974) based on the findings of Imaizumi et al. Comparison of the sequences of the D-loop region with those of others from western Japan in the database showed that *A. speciosus* in Yamaguchi Prefecture did not constitute a distinct group (Fig. S1). These results indicated that *A. speciosus* in Yamaguchi prefecture can be classified into Honshu-Shikoku-Kyushu group.

**Fig. 2.**
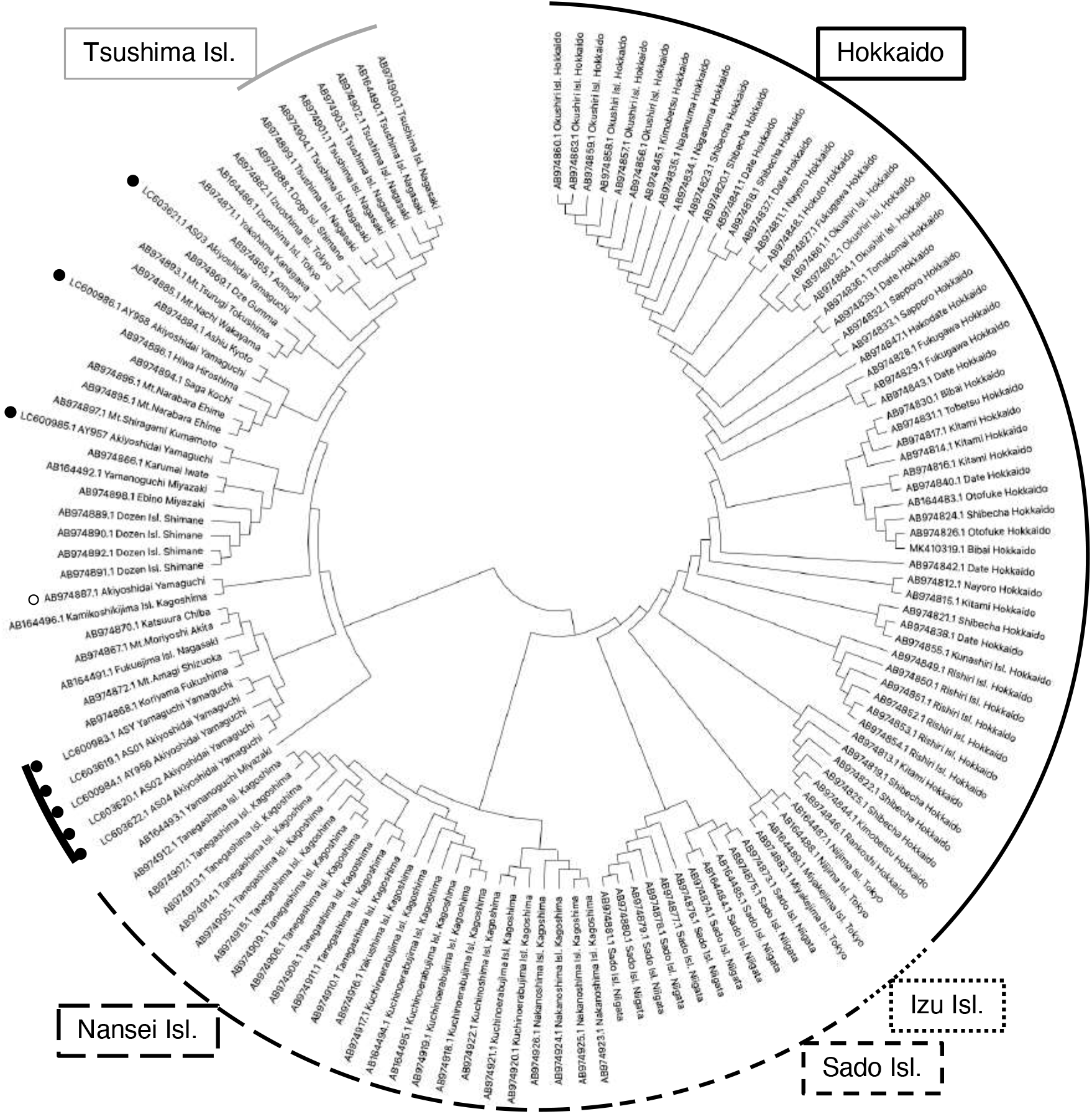
Evolutionary analysis by Maximum Likelihood method. The evolutionary history was inferred by using the Maximum Likelihood method and Tamura-Nei model. The tree with the highest log likelihood is shown. Initial trees for the heuristic search was obtained automatically by applying Neighbor-Join and BioNJ algorithms to a matrix of pairwise distances estimated using the Tamura-Nei model, and then selecting the topology with a superior log likelihood value.

Next, to count the number of chromosomes, we produced cultured cells from the tail tip tissues. As a result, we obtained fibroblast-like cells that migrated and proliferated from the tail tissues (Fig. S2). Chromosomal spreads were prepared by colchicine treatment of these cells (Fig. 3a). We counted the number of chromosomes per cell, which indicated that nearly 80% of the cell had a chromosome number of 2n=46 (Fig. 3b). These results revealed that the chromosome number of *A. speciosus* was 2n=46 in Yamaguchi Prefecture. Previous reports have shown that the number of chromosomes in *A. speciosus* is different between the western and eastern parts of Japan (Shimba and Kobayashi 1969). The number of chromosomes in each region has not been analyzed in detail. Our present study is the first report showing that the number of chromosomes of *A. speciosus* at Akiyoshidai Quasi-National Park in Yamaguchi Prefecture, is 2n=46.

**Fig. 3.**
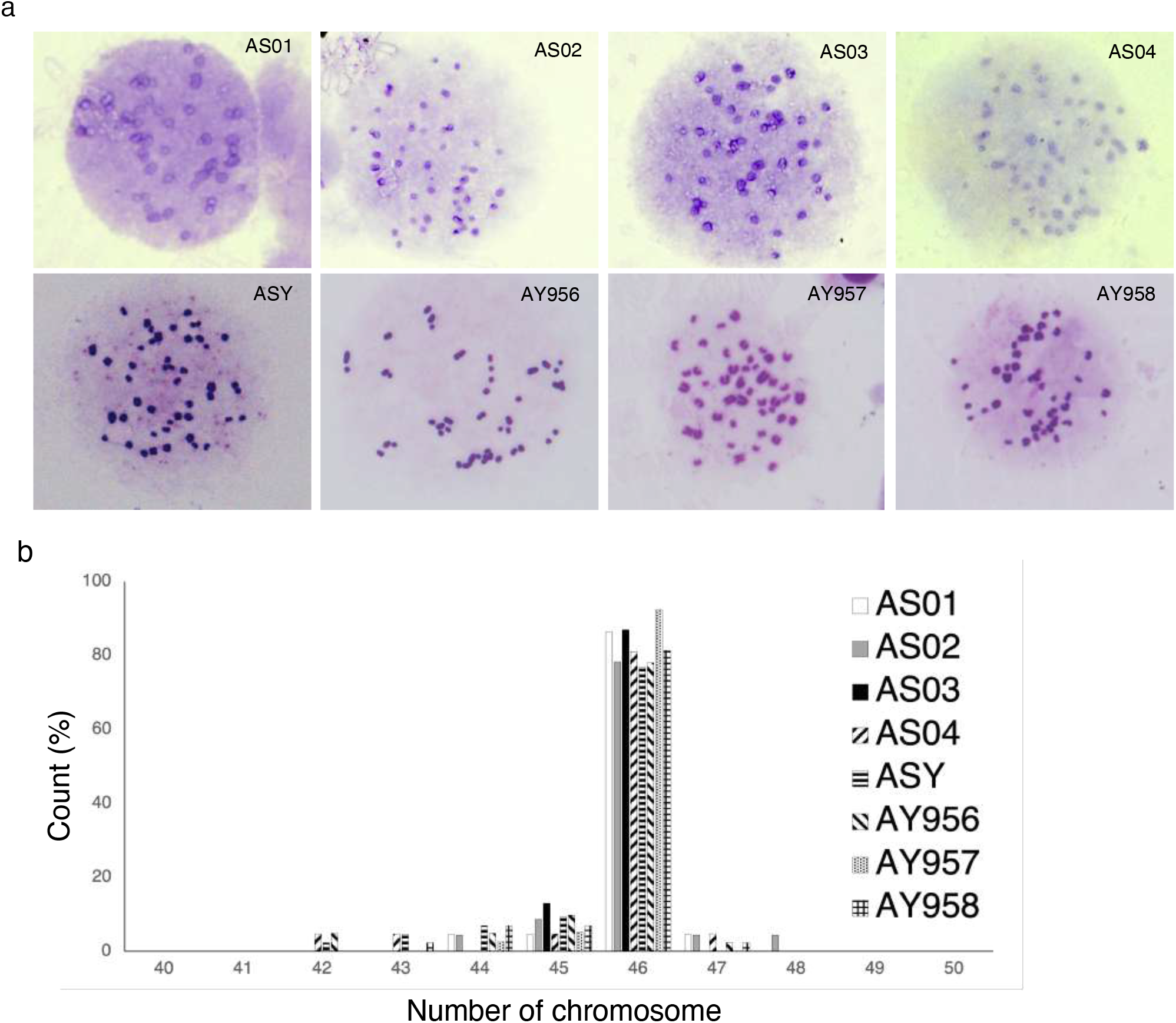
Chromosomal spreads and counts. a) Giemsa stained images of chromosomal spreads of each mice. b) Histogram of the number of chromosomes counted.

In summary, this present study investigated the mtDNA and chromosome number of *A. speciosus* in Yamaguchi Prefecture. Our results showed that *A. speciosus* in Yamaguchi Prefecture had a western Japan-type chromosome number and is classified into a cluster of the Honshu-Shikoku-Kyushu group. Focusing on *A. speciosus* in Yamaguchi Prefecture within Japan as a whole, it is possible that some of *A. speciosus* are genetically clustered. Therefore, it should be necessary to monitor the domestic invasion of *A. speciosus* involving the movement of natural persons and goods. Continuous monitoring of the large Japanese field mice population might be necessary to reveal the dynamics of genetic diversity around Yamaguchi Prefecture.

## Acknowledge

This study was supported by Research and survey program of Akiyoshidai Karst Plateau Academic Center, Yamaguchi University to I.H.

The authors would like to acknowledge the technical expertise of The DNA Core facility of the Center of Gene Research, Yamaguchi University, supported by a grant-in-aid from the Ministry of Education, Science, Sports and Culture of Japan.

---

*Table S1 Primer sequences*

*Table S2 The captured mice and their accession numbers*

**Fig. S1.**
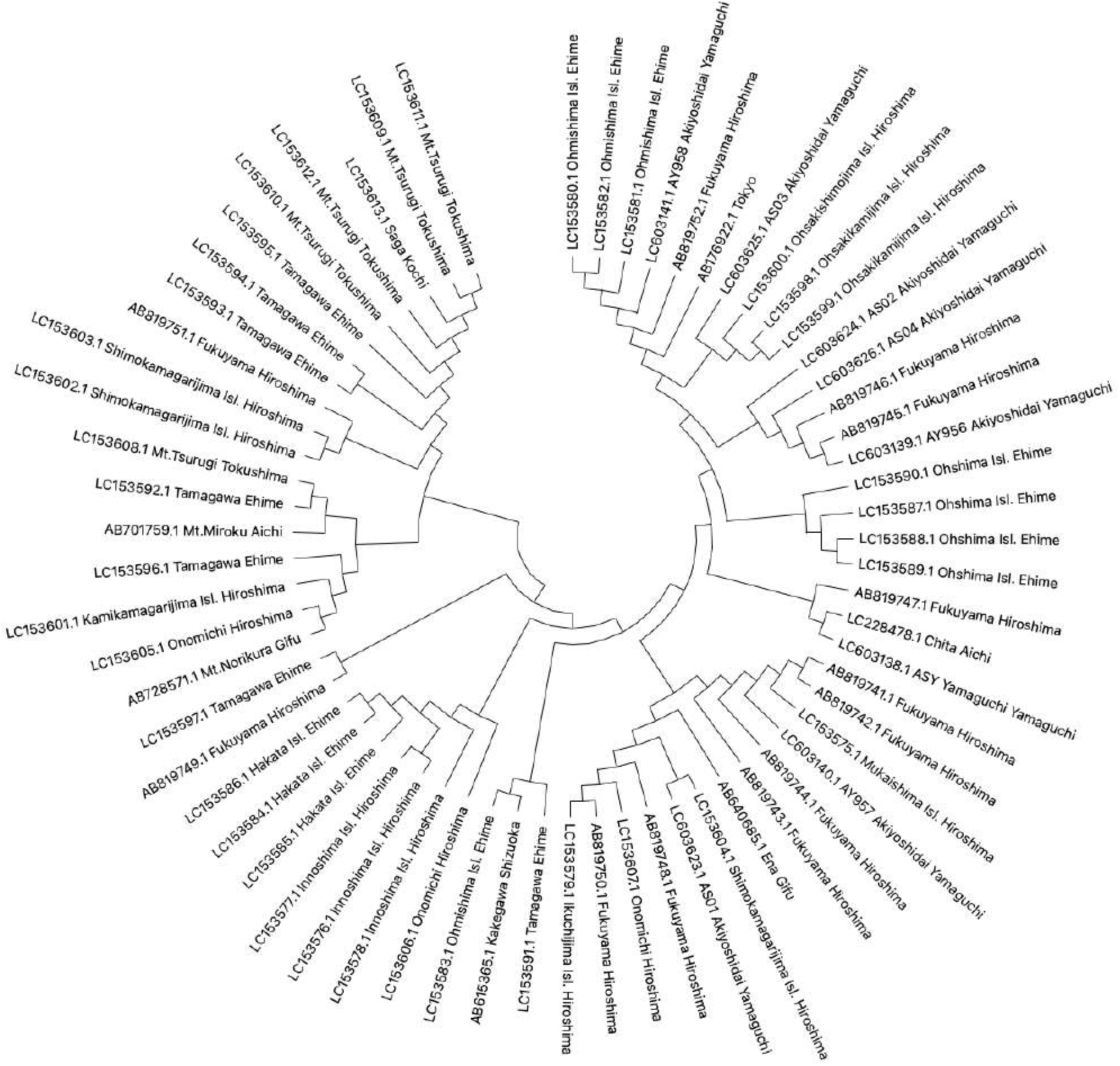
Phylogenetic tree analysis of D-loop region sequences. Maximum Likelihood method was used as in Fig. 2.

**Fig. S2.**
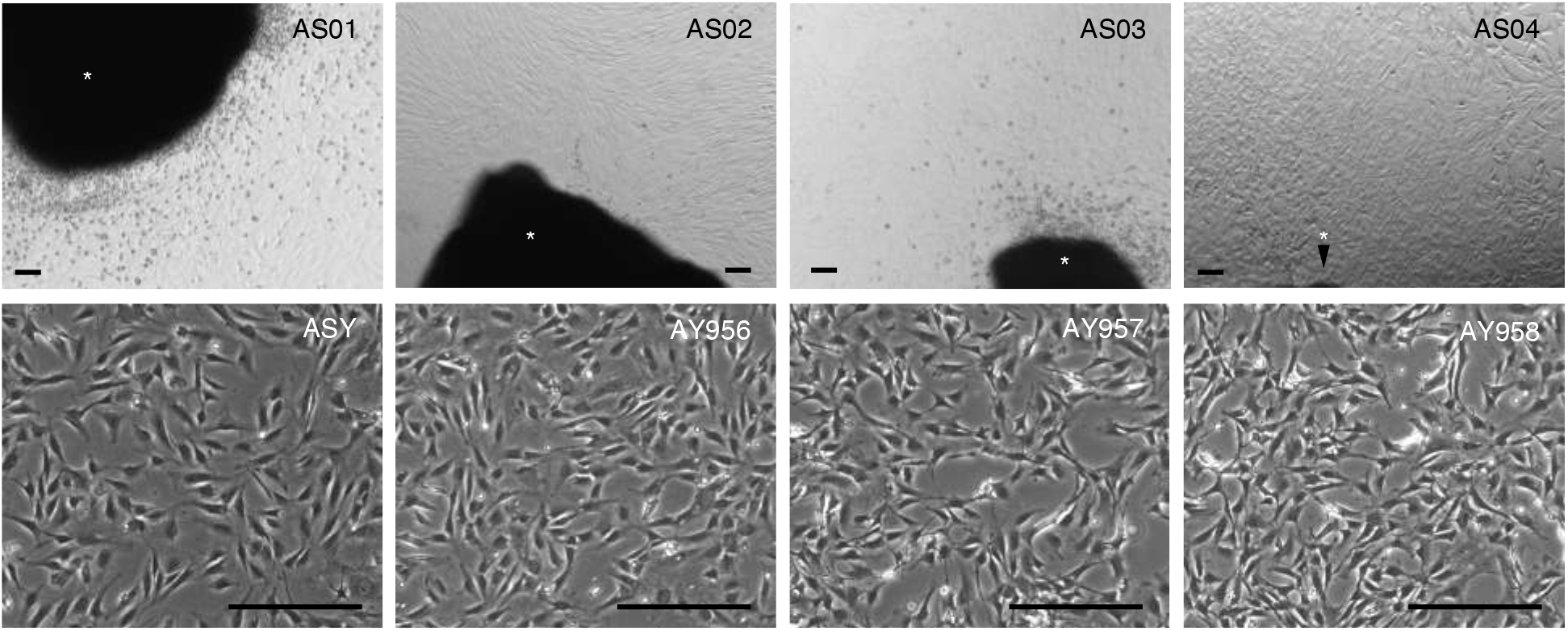
Recovery of cells from tail tip. The upper panels showed the initial stage of culture. The lower showed the cells after passaging. Asterisks indicated tail tips. Bar=100 μm.

## References

Ariyoshi, K., Miura, T., Kasai, K., Goh, V.S.T., Fujishima, Y., Nakata, A., Takahashi, A., Shimizu, Y., Shinoda, H., Yamashiro, H., Seymour, C., Mothersill, C. and Yoshida, M.A. 2020. Environmental radiation on large Japanese field mice in Fukushima reduced colony forming potential in hematopoietic progenitor cells without genomic instability. International Journal of Raddiation Biology 1–12.

Ariyoshi, K., Miura, T., Kasai, K., Akifumi, N., Fujishima, Y. and Yoshida, M.A. 2018. Raddiation-induced bystander effect in large Japanese field mouse (Apodemus speciosus) embryonic cells. Radiation and Environmental Biophysics 57(3): 223–231.

Azuma, R., Hatanaka, Y., Shin, S.W., Murai, H., Miyashita, M., Anzai, M. and Matsumoto, K. 2020. Developmental competence of interspecies cloned embryos produced using cell from large Japanese field mice (Apodemus speciosus) and oocyte from laboratory mice (Mus musculus domesticus). Journal of Reproduction and Development 66(3): 255–263.

Fujishima, Y., Nakata, A., Ujiie, R., Kasai, K., Ariyoshi, K., Goh, V.S.T., Suzuki, K., Tazoe, H., Yamada, M., Yoshida, M.A. and Miura, T. 2020. Assessment of chromosomal aberrations in large Japanese field mice (Apodemus speciosus) In Namie Town, Fukushima. International Journal of Radiation Biology 1–9.

Hanazaki, K., Tomozawa, M., Suzuki, Y., Kinoshita, G., Yamamoto, M., Irino, T. and Suzuki, H. 2017. Estimation of evolution rates of mitochondrial DNA in two Japanese field mouse species based on calibrations with quaternary environmental changes. Zoological Science 34(3): 201–210.

Hirota, T., Hirohata, T., Mashima, H., Satoh, T. and Obata, Y. 2004. Population structure of the large Japanese field mouse, Apodemus speciosus (Rodentia: Muridae), in suburban landscape, based on mitochondrial D-loop sequences. Molecular Ecology 13(11): 3275–3282.

Kawagoshi, T., Shiomi, N., Takahashi, H., Watanabe, Y., Fuma, S., Doi, K., Kawaguchi, I., Aoki, M., Kubota, M., Furuhata, Y., Shigemura, Y., Mizoguchi, M., Yamada, F., Tomozawa, M., Sakamoto, S.H., Yoshida, S. and Kubota, Y. 2017. Chromosomal aberrations in large Japanese field mice (Apodemus speciosus) captured near Fukushima Dai-Ichi nuclear power plant. Environmental Science and Technology 51(8): 4632–4641.

Kumar, S., Stecher, G., Ki, M., Knyaz, C. and Tamura, K. 2018. MEGA X: Molecular Evolution Genetics Analysis across computing platforms. Molecular Biology and Evolution 35(6): 1547–1549.

Kuwahara, S., Mizukami, T., Omura, M., Hagihara, M., Iinuma, Y., Shimizu, Y., Tamada, H., Tsukamoto, Y., Nishida, T. and Sasaki, F. 2000. Seasonal changes in the hypothalamo-pituitary-testes axis of the Japanese wood mouse (Apodemus speciosus). The anatomical Record 260(4): 366–372.

Matsubara, K., Nishida-Umehara, C., Tsuchiya, K., Nukaya, D. and Matsuda, Y. 2004. Karyotypic evolution of Apodemus (Muridae, Rodentia) inderred from comparative FISH analysis. Chromosome Research 12(4): 383–395.

Matsunami, M., Endo, D., Saitou, N., Suzuki, H. and Onuma, M. 2017. Draft genome sequence of Japanese wood mouse, Apodemus speciosus. Data Brief 16: 43–46.

Meguro, K., Kamotsu, K., Ohdaira, T., Nakagata, N., Nakata, A., Fukumoto, M., Miura, T. and Yamashiro, H. 2019. Induction of superovulation using inhibin antiserum and competence of embryo development in wild large Japanese field mice (Apodemus speciosus). Reproduction in Domestic Animals 54(12): 1637–1642.

Okano, T., Ishiniwa, H., Onuma, M., Shindo, J., Yokohata, Y. and Tamaoki, M. 2016. Effects of environmental radiation on testes and spermatogenesis in wild large Japanese field mice (Apodemus speciosus) from Fukushima. Scientific Reports 6: 23601.

Okano, T., Onuma, M., Ishikawa, H., Azuma, N., Tamaoki, M., Nakajima, N., Shindo, J. and Yokohata, Y. 2015. Classification of the spermatogenic cycle, seasonal changes of seminiferous tubule morphology and estimation of the breeding season of the large Japanese field mouse (Apodemus Speciosus) in Toyama and Aomori Prefectures, Japan. Journal of Veterinary Medical Science 77(7): 799–807.

Sakai, Y., Sakamoto, S.H., Kato, G.A., Iwamoto, N., Ozaki, R., Eto, T., Shinohara, A., Morita, T. and Koshimoto, C. 2013. Rearing method to induce natural mating of the large Japanese field mouse, Apodemus speciosus. Honyuruikagaku 53(1): 57–65. (In Japanese, Abstract in English)

Sato, J.J., Tasaka, Y., Tasaka, R., Gunji, K., Yamamoto, Y., Takada, Y., Uematsu, Y., Sakai, E., Tateishi, T. and Yamaguchi, Y. 2017. Effect of isolation by continental islands in the Seto inland sea, Japan, on genetic diversity of the large Japanese field mouse, Apodemus Speciosus (Rodentia: Muridae), inferred from the mitochondrial Dloop region. Zoological Science 34(2): 112–121.

Shimba, H. and Kobayashi, T. 1969. A Robertsonian type polymorphism of the chromosomes in the field mouse, Apodemus speciosus. Japanese Journal of Genetics 44(3):117–122.

Stecher, G., Tamura, K. and Kumar, S. 2020. Molecular Evolutionary Genetics Analysis (MEGA) for macOS. Molecular Biology and Evolution 37(4): 1237–1239.

Suzuki, Y., Tomozawa, M., Koizumi, Y., Tsuchiya, K. and Suzuki, H. 2015. Estimating the molecular evolution rates of mitochondrial genes referring to quaternary ice age events with inferred population expansion and dispersals in Japanese Apodemus. BMC Evolutionary Biolology 15: 187.

Takino, S., Yamashiro, H., Sugano, Y., Fujishima, Y., Nakata, A., Kasai, K., Hayashi, G., Urushihara, Y., Suzuki, M., Shinoda, H., Miura, T. and Fukumoto, M. 2017. Analysis of the effect of chronic and low-dose radiation exposure on spermatogenic cells of male large Japanese field mice (Apodemus Speciosus) after the Fukushima Daiichi nuclear power plant accident. Radiation Research 187(2): 161–168.

Tamura, K., and Nei, M. 1993. Estimation of the number of nucleotide substitutions in the control region of mitochondrial DNA in humans and chimpanzees. Molecular Biology and Evolution 10(3): 512–526.

Tsuchiya, K. 1974. Cytological and biochemical studies of Apodemus speciosus group in Japan. The Journal of Mammalogical Society of Japan 6(2):67–87. (In Japanese, Abstract in English)

Yamagishi, M., Matsubara, K. and Kasaizumi, M. 2012. Molecular cytogenetic identification and characterization of Robertsonian chromosomes in the large Japanese field mouse (Apodemus speciosus) using FISH. Zoological Science 29(10): 709–713.

