## Supplementary Table for "Genetic characteristics of *Apodemus speciosus* at Akiyoshidai Quasi-National Park in Yamaguchi Prefecture"

| Name | Sequences (5'->3') | Anealing Temperature | Discription |
|---|---|---|---|
| L14724 | CGAAGCTTGATATGAAAAACCATCGTTG | 50 | Cytb |
| H15915 | AACTGCAGTCATCTCCGGTTTACAAGAC | | *1 |
| R-L14724 | CAGGAAACAGCTATGACCGATATGAAAAACCATCGTTG | 50 | Cytb |
| SNH655 | TGTAAAACGACGGCCAGTTGTGTAGTATGGGTGGAATGG | | *1 |
| SNL497 | CAGGAAACAGCTATGACCCCTAGTAGAATGAATCTGAGG | 50 | Cytb |
| R-H15916 | TGTAAAACGACGGCCAGTGTCATCTCCGGTTTACAAGA | | *1 |
| M15997 | TCCCCACCATCAGCACCCAAAGC | 63 | D-loop |
| H16401 | TGGGCGGGTTGTTGGTTTCACGG | | *2 |

*1: Suzuki et al. 2015, *2: Hirota et al. 2004.

Table S1 Primer sequences

| | Accession number | |
|---|---|---|
| | Cytb | D-loop |
| ASY | LC603138 | LC603138 |
| AY956 | LC603139 | LC603139 |
| AY957 | LC603140 | LC603140 |
| AY958 | LC603141 | LC603141 |
| AS01 | LC603619 | LC603623 |
| AS02 | LC603620 | LC603624 |
| AS03 | LC603621 | LC603625 |
| AS04 | LC603622 | LC603626 |

Table 2 The captured mice and their accession numbers.
